# Response of Various Concentrations Of Indole Acetic Acid (IAA) to Morphological Attributes in Different Plum (*Prunus domestica* L.) Curltivars

**DOI:** 10.1101/2024.09.19.614029

**Authors:** Kaleem Ullah, Samin Jan, Roshan Zamir, Abdul Majeed, Bakhtiyar Ahmad, Gouhar Rehman

## Abstract

This study was conducted to evaluate the effects of indole-3-acetic acid (IAA) on budwood treated with different concentrations of indole acetic acid i.e. ((300, 600, 900 and 1200 ppm) of three plum varieties (Fazle Manani, Santa Rosa and Red Beauty) in respect of various morphological parameters e.g. days to sprouting, plant height, number of branches, percent sprouting, number of leaves, budling diameter and internode length. Results revealed that different varieties of plum had different responses to the applied concentrations in different plum cultivars. Cultivar Fazle manani showed significant decline in days to sprouting (at 300 ppm dose), plant height (at 600 dose), percent sprouting and budling diameter (at all concentrations) while increase in number of branches (1200 dose), number of leaves (all concentrations except 600 ppm), and internode length (at 300 ppm and 900 ppm concentrations). Days to sprouting in cultivar Santa rosa were non-significantly affected while plant height and internode length were significantly reduced at all concentrations particularly at 300 ppm. Number of branches, number of leaves and percent sprouting increased significantly at 300 ppm and 900 ppm concentrations. Cultivar red beauty exhibited significant decline in days to sprouting at 600 ppm and 900 ppm concentrations while elevated plant height at 900 concentrations. Number of branches, percent sprouting, number of leaves and budling diameter in Red Beauty were inhibited by all concentrations. However, internode length was increased significantly at 300 ppm, 600 ppm and 1200 ppm concentrations but decreased at 900 ppm concentrations. The study revealed that the most effective dose was 300 ppm. These findings demonstrate the differential sensitivity of plum cultivars to IAA and highlight its potential for regulating their growth and morphology.

## 1. INTRODUCTION

Plant growth regulators (PGRs) are chemical substances which have regulatory effects on different aspects of plants such as root and shoot growth, fruit development & regulation and other developmental processes (Fishel, 2006). Five major classes of plant growth regulators have been identified which are auxins, gibberellins, cytokinins, ethylene and abscisic acid that are produced in different locations of plants under different developmental phases (Small and Degenhardt, 2018). The Plant growth regulators (PGRs) have been recognized for their effects on cells (cell division, expansion, and differentiation) regulation of physiological processes, and developmental process such as formation of roots, shoots, buds, flowers, and fruits (Flasiński and Hąc-Wydro, 2014). Auxin (indole-3-acetic acid) which has complex biosynthetic pathways in plants induces growth and development of plants through diverse mechanisms (Zhao, 2010).

Plum (*Prunus domestica* L.) is the most important stone fruit in Pakistan next to peaches (Westwood, 1993). Budding of European plum (*P. domestica*) mostly takes place in plum, peach, almond and apricot as root stock.

In Pakistan plum is cultivated on large acreage (more than 7000 hectares) and is the world most growing country in the world, and it rank 17^th^ in production and its total production is 67000 tones. The most growing and most productive provinces of Pakistan are Baluchistan and Khyber Pakhtunkhwa. In Baluchistan Kalat, Mastung, Pashin and Quetta while in khyber pakhtunkhwa Mardan, Nowshera and Swat are the most producing areas of the mention provinces. Khyber Pakhtunkhwa share 47% of the total production of the country. While share of the district Swat is 17% out of the total production of the province (GOP, 2009).

The focus is on traditional breeding techniques to develop plum cultivars with high yields and quality of fruits, resistance to biotic and abiotic constraints, and better adaptability to a diverse range of climatic conditions (Okie and Hancock, 2008). However, traditional breeding strategies are laborious and time consuming and thus, novel methods must be employed to minimize the issues related to traditional breeding. In recent years, genetic engineering and biotechnological interventions have been linked with enhancing the growth, yield, and productivity of plum (Topp *et al*., 2012). According to Neumüller (2011) genetic engineering and the use of markers are important tools for improving plums; however, such techniques are limited to the embryonic transformation.

Asexual propagation of plum through budding, cuttings and grafting has been shown an effective mean to overcome the problems of climatic variability, tolerance to soil, pathogenic stresses, and yield efficiency (Wolella, 2017). Using different rootstock for budding, grafting and cuttings can lead to increased genetic diversity, efficient adaptability, and improved propagation of plums. Moreover, propagation of different cultivars which have overlapping flowering period is a good strategy to overcome the problems of self-incompatibility (Ohata *et al*., 2017).

Plum is an important stone fruit propagated by budding on rootstocks have many biotic and abiotic constraints. The study was conducted to find out the effect of indole -3-acetic acid (IAA) to buds wood and responses of three plum cultivars (Fazle manani, Santa rosa and Red beauty) in respect of various morphological characters.

## 2. METHODOLOGY

The research was carried out at the Nuclear Institute for Food and Agriculture (NIFA) Peshawar in October 2022. Three plum varieties i.e. Fazle manani, Santa rosa and Red beauty, were chosen as scion cultivars to investigate the effects of different concentrations of indole -3-aceticacid (IAA) on bud development at various growth stages to check the morphological responses of various plum cultivars. The scion varieties were grafted onto Mariana plum cuttings serving as the rootstock in the experimental field at NIFA, Peshawar. The experimental design followed was Randomized Complete Block Design (RCBD) with two main factors: Varieties and Treatments. The experiment was replicated three times, and each replication consisted of 5 plants per treatment.

### 2.1. Treatment of buds with plant growth regulators (PGRs)

Buds wood was collected from three cultivars of plum (Fazli manani, Santa Rosa, and Red Beauty) trees grown at NIFA Peshawar orchard during the first week of October 2022. The collected buds were brought to the laboratory and thoroughly washed under running clean tap water. Different concentrations (viz 300, 600, 900 and 1200 ppm) of indole acetic acid (IAA) were prepared along with control (Water without IAA) and stored in sterilized flasks (volume 1 liter) in a refrigerator at a temperature (20^°^C to 25 ^°^C) in the tissue culture laboratory of NIFA Peshawar. The collected buds were soaked separately in 04 different concentrations of indole acetic acid (IAA) for 12 hours. One set consists of three (03) different varieties of plum. bud wood was used as control for budding on rootstocks.

Mariana plum cuttings of a local cultivar of plum were planted in February 2022. Bud woods were collected from three plum cultivars and treated with different concentrations of indole acetic acid (IAA) were budded into Mariana cuttings using T-budding techniques. The budding process was carried out during mid-October 2022 while data for buds sprouting and other morphological tributes were collected during April-Sep. 2023. For each treatment and cultivar, 15 Mariana cuttings were used.

### 2.2. Ethical Statement

The research conducted on the plants strictly followed established national and international procedures and guidelines. This adherence ensured that all experimental methods were ethically sound and scientifically valid.

### 2.3. Experimental design

The research utilized two factorials completely randomized blocked designs (RCBD) with three replications. The initial factor involved the collection of buds from three different plum cultivars i.e. Fazli manani, Santa rosa, and Red beauty. The second factor entailed the use of diverse concentrations of Indole Acetic Acid (IAA). Each treatment underwent three replications, and within each replication, there were 5 plants per treatment. The Mariana cuttings employed in the experiment were sourced from a local plum cultivar.

### 2.4. Data Analysis

The collected data were subjected to analysis of variance (ANOVA) at p≤0.05. A two-factor ANOVA was used to estimate the effect of two independent variables on dependent variables. Independent variables of this study were:

a. Different concentrations of Indole acetic acid (IAA)
b. Different cultivars of plums

While dependent variables were the growth parameters.

## 3. RESULTS

### 3.1. DAYS TO SPROUTING

Response of different varieties of plum to different concentrations of IAA for days to sprouting is presented in Fig.1. The number of days to sprouting was investigated both in control and treated budwood of three Cultivars of plum i.e Red beauty Santa Rosa and Fazli manani. It was observed that cultivar Fazle manani exhibited maximum days to sprouting (137 days) with IAA at controlled, followed by 900, 1200, and 600 concentrations where it took (125), (122) and (120.67) days respectively. The lowest days to sprouting were observed at 300 ppm concentration, dose of the tested PGR which reflected its efficacy in reducing the days to sprouting.

**Fig 1.**
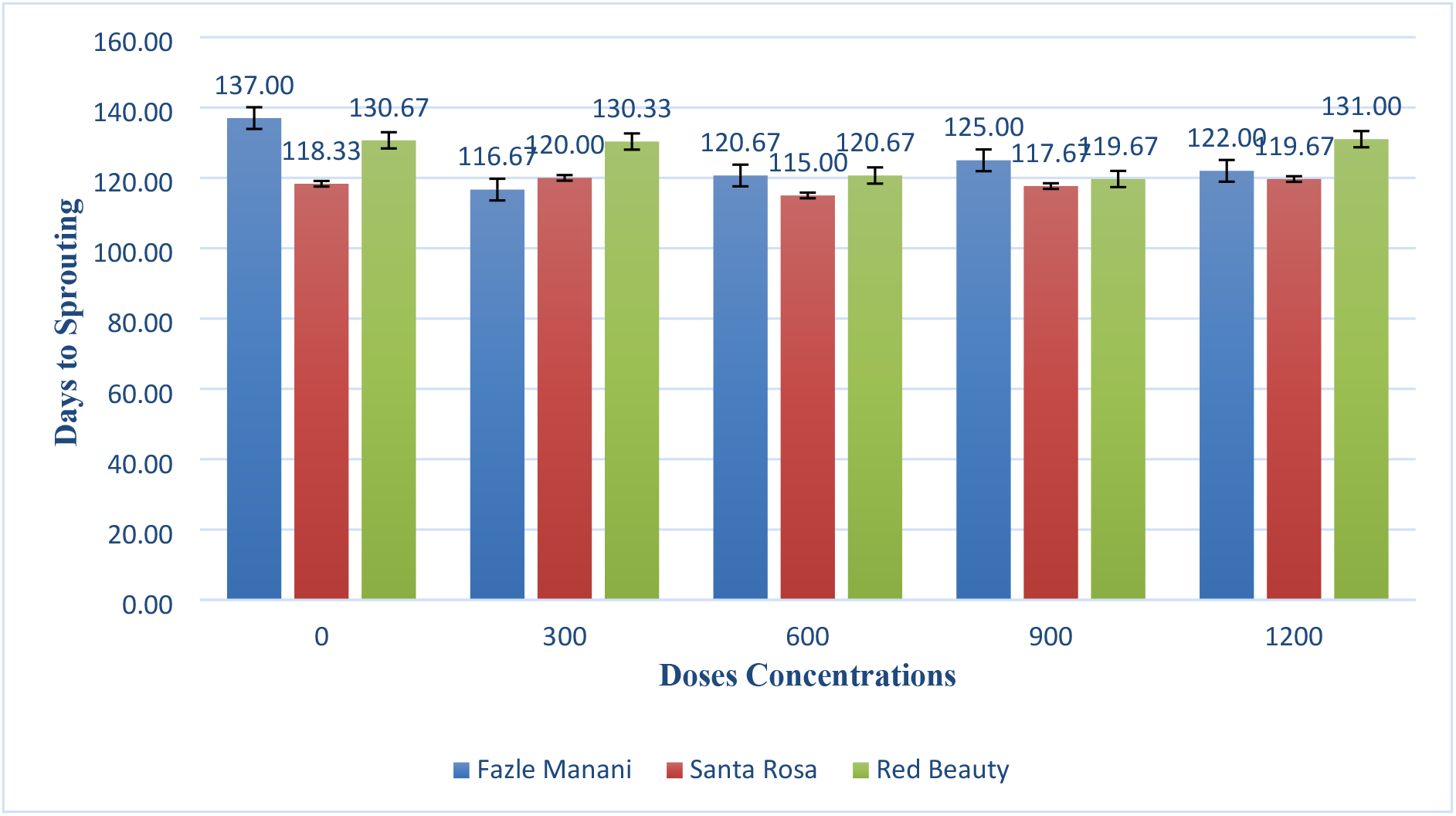
Responses of plum varieties to various concentration of IAA in respect of days to sprout of buddling.

In case of cultivar Santa Rosa, variable responses were observed against different concentrations of the applied PGR. However, maximum days to sprouting (120) days were observed at 300 concentrations followed by (119.67), (118.33) and (117.67) days at 1200, 0 (control) and 900 concentrations respectively. Indole acetic acid (IAA) having 600 concentration was found to be the most suitable in reducing days to sprouting to (115.00) days as compared to control and other applied concentrations.

In Red Beauty, the cultivar response to 0 and 300 concentrations was almost similar where (130.67) and (130.33) days to sprouting were observed. This was followed by a slight decline in days to sprouting (131) days at 1200 dose. The most effective concentrations were recorded as 900 and 600 which reduced days to sprouting to (119.67) and (120.67) days respectively. Data revealed that cultivars showed different responses to different concentrations of IAA.

### 3.2. EFFECTS OF IAA ON PLANT HEIGHT (Cm)

Data for plant height of three cultivars of plum in response to different concentrations of IAA is shown in Fig.2. Data revealed that maximum plant height (99.67 cm) was observed in *Fazli manani* at 300 concentrations which was followed by (99.00 cm) at controlled. Plant height was significantly reduced to (69.9 cm) at 600 concentrations. At 900 and 1200 concentrations, plant height was observed (82.67 cm) and (79.5 cm) respectively.

**Fig 2.**
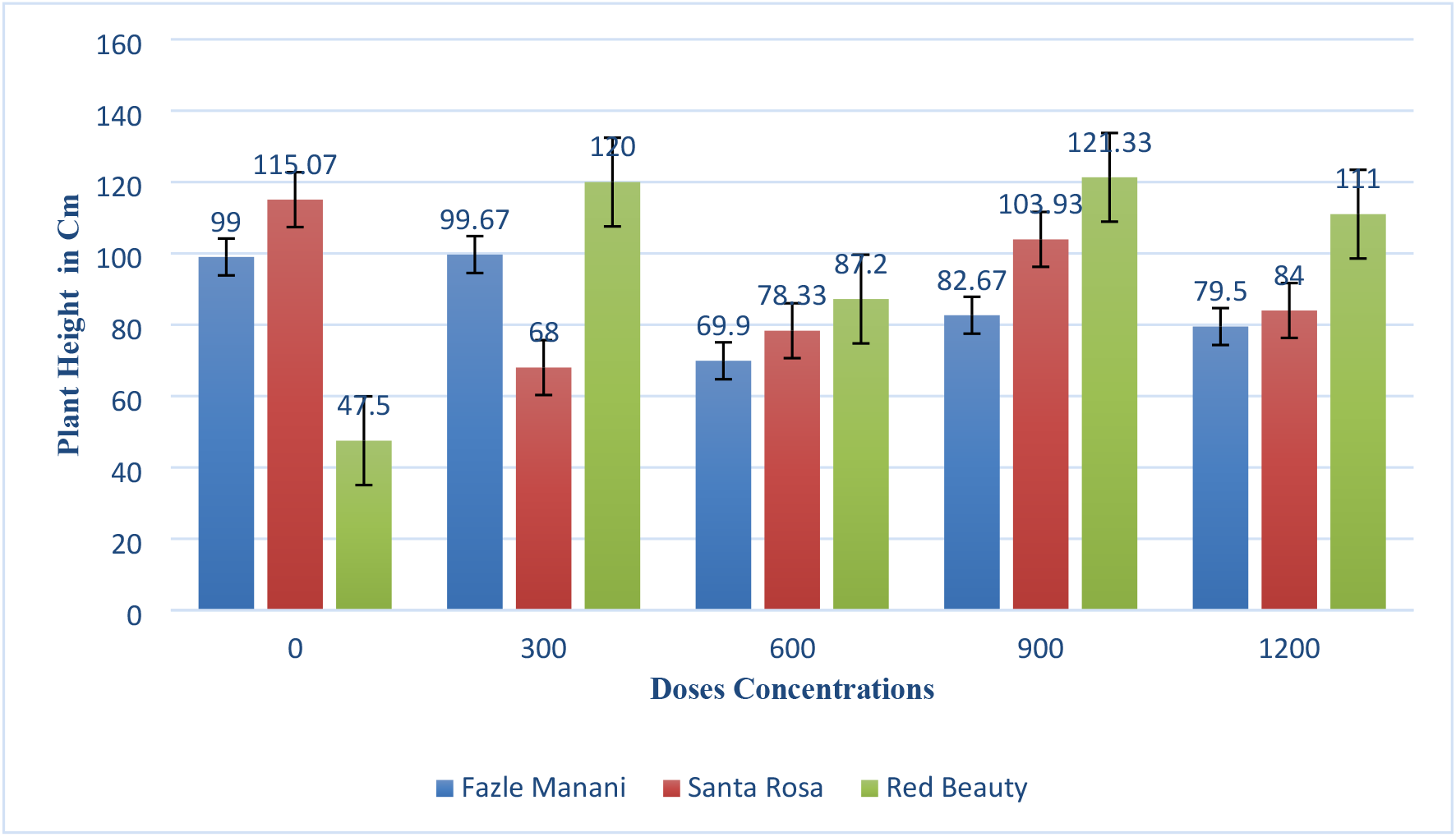
Response of different plum cultivar to various concentrations of IAA in respect of plant height in cm

In case of *Santa rosa* maximum plant height (115.00 cm) occurred in controlled, while the lowest (68.00 cm) was observed in 300 concentrations. At concentrations 600, 900 and 1200 plant height was recorded (78.33), (103.99) and (84.00 cm) respectively. As compared to control all the concentrations have negative effect on plant height. This inhibitory effect in santa rosa might be due to genetic makeup or changes in physiology of the said variety.

Like other two varieties red beauty maximum plant height (121.33) was observed under the influence of 900 concentrations, which was followed by (120.00 cm) at 300 concentrations. The lowest plant height (47.50 cm) was recorded at controlled. Other concentrations 600 and 1200 also had a stimulatory effect on plant height recorded as (87.20) and (111.00 cm) respectively.

### 3.3. EFFECTS OF IAA ON NUMBER OF BRANCHES PER BUDDLING

Response of the three different cultivars of plum for various concentrations of IAA has been observed for number of branches as shown in Fig.3. The data revealed that highest number of branches per buddling (7.58) was recorded in Fazli manani at concentrations 1200, followed by (6.63) at 600 concentrations. The lowest number of branches (4.67) was recorded at controlled concentrations. The other concentrations 300 and 900 also had stimulatory effects on number of branches recorded as (5.00) and 5.55 respectively.

**Fig 3.**
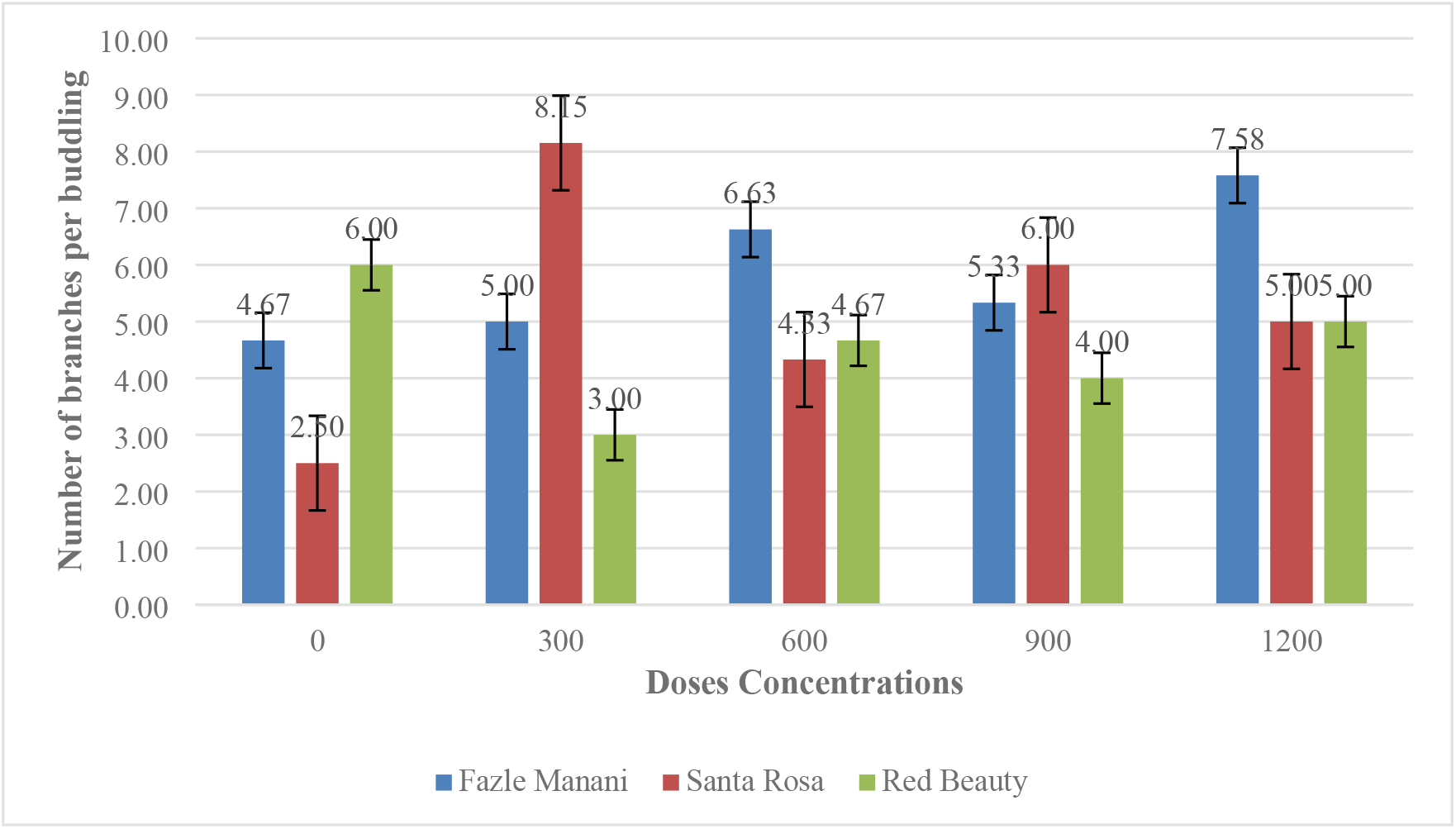
Responses of plum cultivar against various concentrations of IAA in respect of number of branches per buddling

In case of Santa rosa the maximum number of branches (8.15) was observed at 300 concentrations, followed by (6.00) at 900 concentrations. The lowest number of branches (2.50) was recorded at controlled. The other concentrations 600 and 1200 also have some stimulatory effect and were recorded as (4.33) and (5.00) respectively.

Like other two varieties of plum, in red beauty maximum number of branches (6.00) per buddling had observed at controlled concentrations, followed by (5.00) numbers of branches at 1200 concentrations. The lowest number of branches (3.00) had observed at 300 concentrations. The other concentrations have some inhibitory effect on number of branches and recorded as (4.67) and (4.00) at 600 and 900 concentrations respectively.

### 3.4. EFFECTS OF IAA ON PERCENT SPROUTING

The varietal responses of different cultivars against different concentrations of IAA have been observed as shown in Fig.4. Maximum percent sprouting 70% was observed in Fazli manani at controlled concentrations, followed by (50.33%) at 300 concentrations. The lowest percent sprouting (47.00%) was observed at 900 ppm concentrations. The other concentrations 600 and 1200 also have stimulatory effect on Percent sprouting and revealed (48.33%) and (48.00%) respectively.

**Fig 4.**
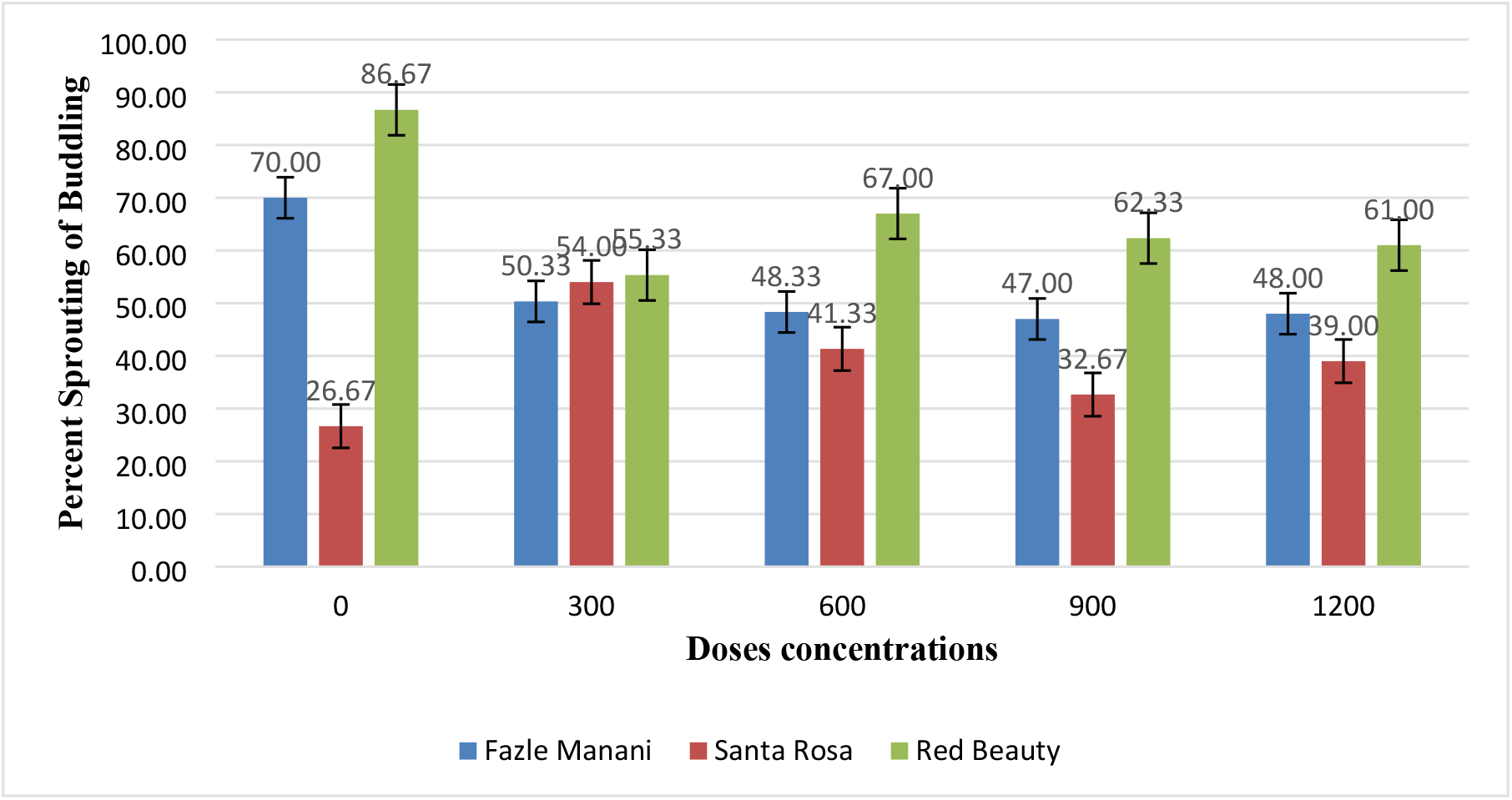
Response of plum cultivar to various concentrations of IAA in respect of percent sprouting of budling

In other plum cultivar santa rosa maximum percent sprouting (54.00%) was occurred at 300 concentrations, followed by (41.33%) at 600 ppm concentrations. The lowest percent sprouting (26.67%) was recorded in controlled treatments. The other concentrations 600 ppm and 1200 ppm has some stimulatory effect on percent sprouting and recorded as (41.33%) and (39.00%) respectively.

Like other two varieties of plum, in red beauty maximum percent sprouting (86.67%) has been observed at controlled concentrations, followed by (67.00%) at 600 concentrations. The lowest percent sprouting 55.00% was observed at 300 concentrations. The other concentrations 900 and 1200 concentrations also had some inhibitory effect on percent sprouting observed was (62.33%) and (61.00%) respectively.

### 3.5. EFFECTS OF IAA ON NUMBER OF LEAVES PER BUDDLING

Responses of three different cultivars of plum have been observed for different concentrations of IAA for number of leaves per budling as shown in Fig.5. The highest number of leaves per buddling (124.00) was observed in Fazli manani at 900 concentrations, followed by (112.33) at 1200 concentrations. The lowest number of leaves per buddling (93.67) was s observed in 600 ppm concentrations. The other dose 300 ppm also had stimulatory effect and (111.00) numbers of leaves per buddling have been recorded.

**Fig 5.**
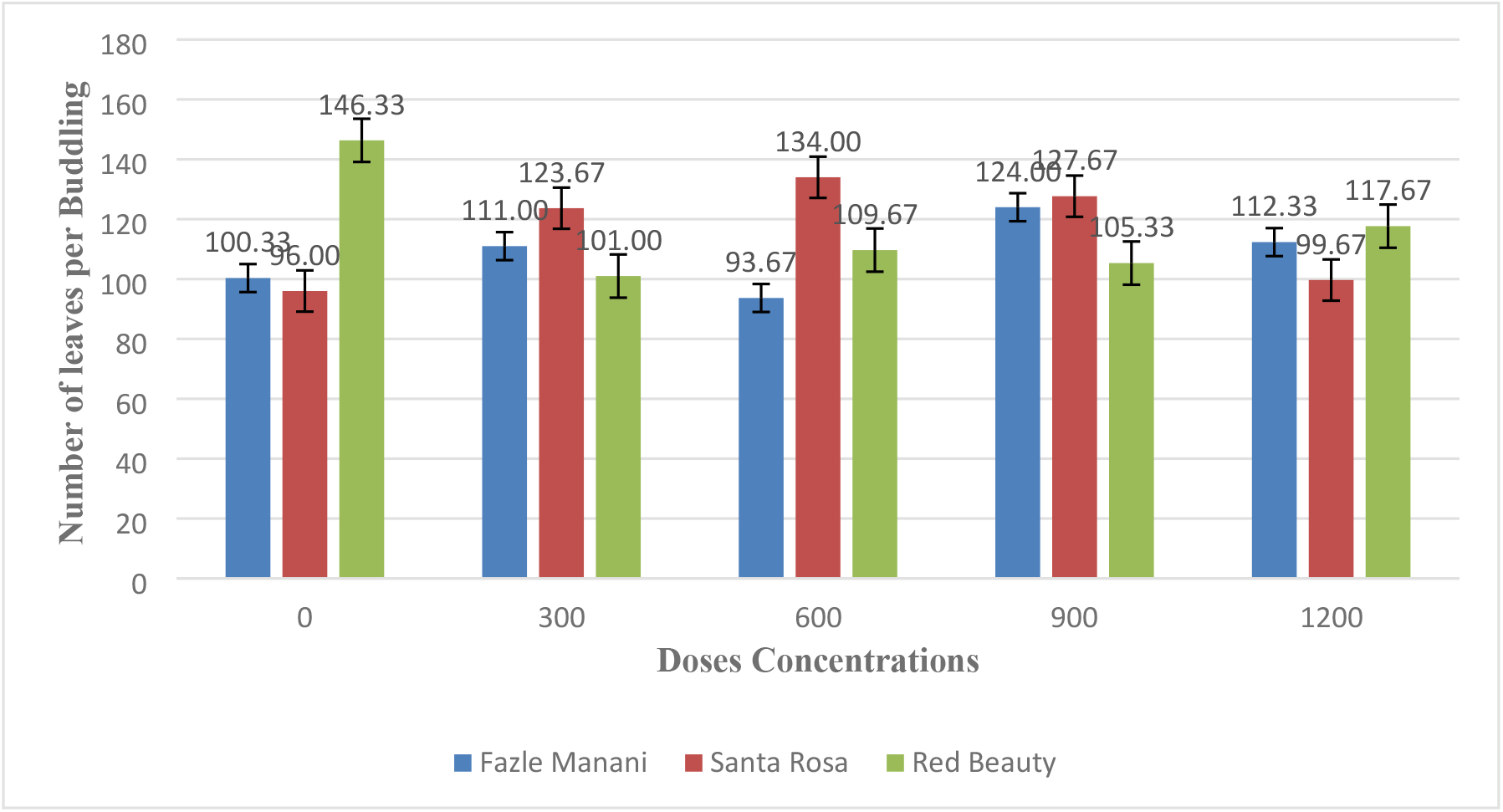
Responses of plum cultivar to various concentrations of IAA in respect of number of leaves per budling

In case of Santa rosa the higher number of leaves per buddling (134.00) was observed at control 600 concentrations, followed by (127.67) at 900 ppm concentrations. The lowest number of leaves per buddling 96.00 was observed at control concentrations. The other concentrations 300 and 1200 also have some stimulatory effect and (123.67) and (99.67) number of leaves per buddling has been recorded respectively.

Like other varieties of plum in red beauty maximum number of leaves per buddling 146.33 has been observed in control concentrations. The other concentrations have inhibitory effect on number of leaves per buddling. The most retarded effect was observed on number of leaves (101.00) at 300 ppm concentrations, followed by 900 ppm, 600 ppm and 1200 ppm concentrations, the number of leaves per buddling were (105.33), (109.67) and (117.67) respectively.

### 3.6. EFFECTS OF IAA ON BUDDLING DIAMETER

The effect of different concentrations of IAA was observed on different varieties of plum in respect of buddling diameter as outlined in Fig.6. In fazle manani maximum buddling diameter (7.78 mm) was recorded in control, while the other treatment has slightly inhibitory effect on the studies parameter. The lowest buddling diameter (6.00 mm) was recorded as at 900 ppm concentrations.

**Fig 6.**
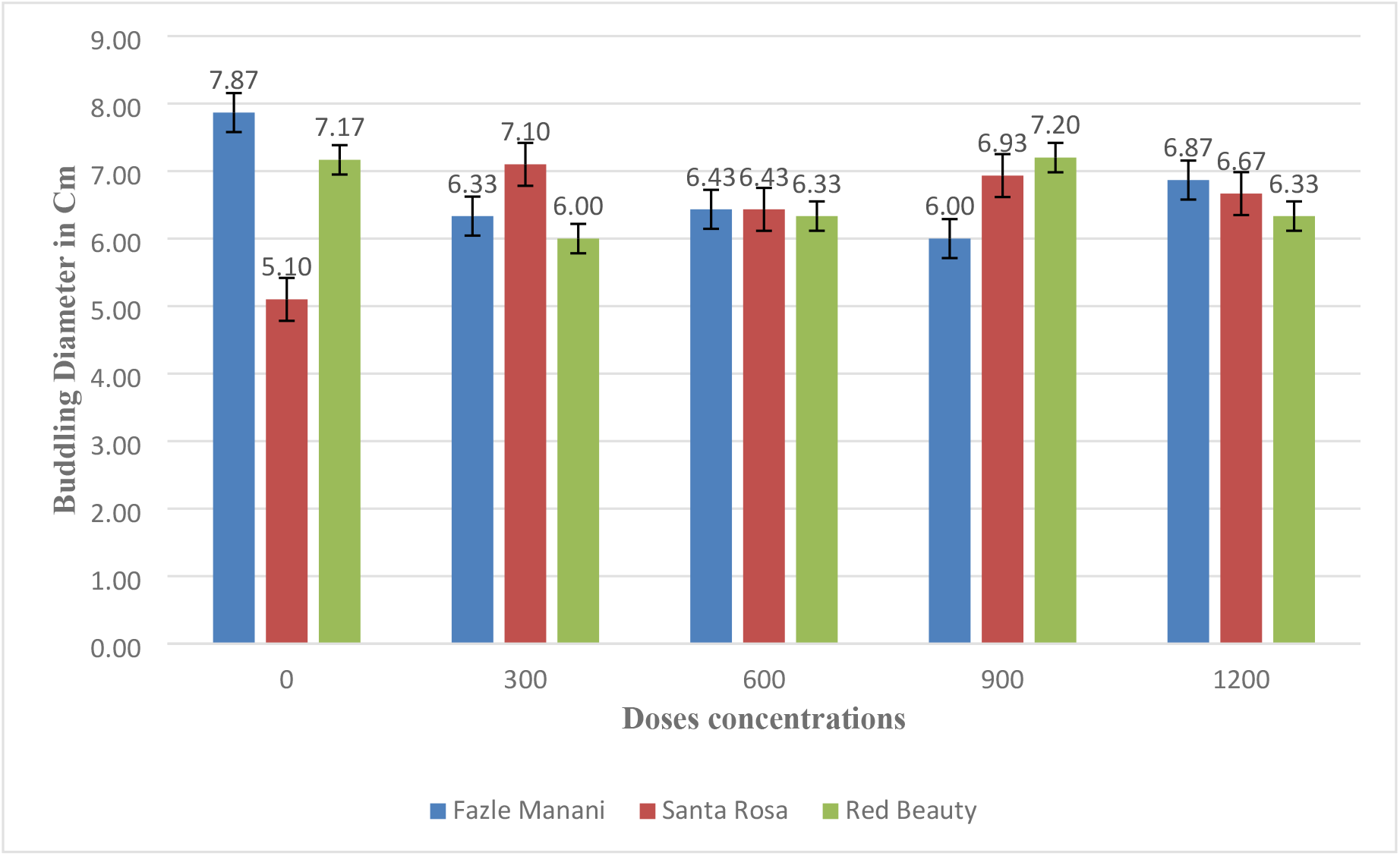
Responses of plum cultivar to various concentrations of IAA in respect of buddling diameter

In case of plum cultivar Santa rosa the highest buddling diameter (7.10 mm) was observed in 300 concentrations, followed by (6.93 mm) at 900 concentrations. The lowest buddling diameter was observed at control concentrations. The other concentrations have stimulatory effect at 600 and 900 concentrations the buddling diameter was observed (6.43 cm) and (6.67 mm) respectively.

In cultivar red beauty the highest buddling diameter (7.20 mm) was observed at 900 concentrations, followed by (7.17 mm) at control. The other concentrations 300 ppm, 600 ppm, and 1200 ppm have slightly inhibitory effect on the buddling diameter revealing (6.00 mm), (6.33 cm), and (6.33 mm) respectively.

### 3.7. EFFECTS OF IAA ON INTERNODE LENGTH

The effect of different concentrations of IAA on buds and responses of three different cultivar of plum in respect of internode length were observed in Fig.7. In Fazle manani the highest internode length (1.97 cm) was observed at 300 concentrations, followed by (1.90 cm) at 900 concentrations. The lowest internode length (1.77 cm) was observed at control and 600 concentrations. The other concentrations 1200 had a stimulatory effect on internode length revealed as (1.80 cm).

**Fig 7.**
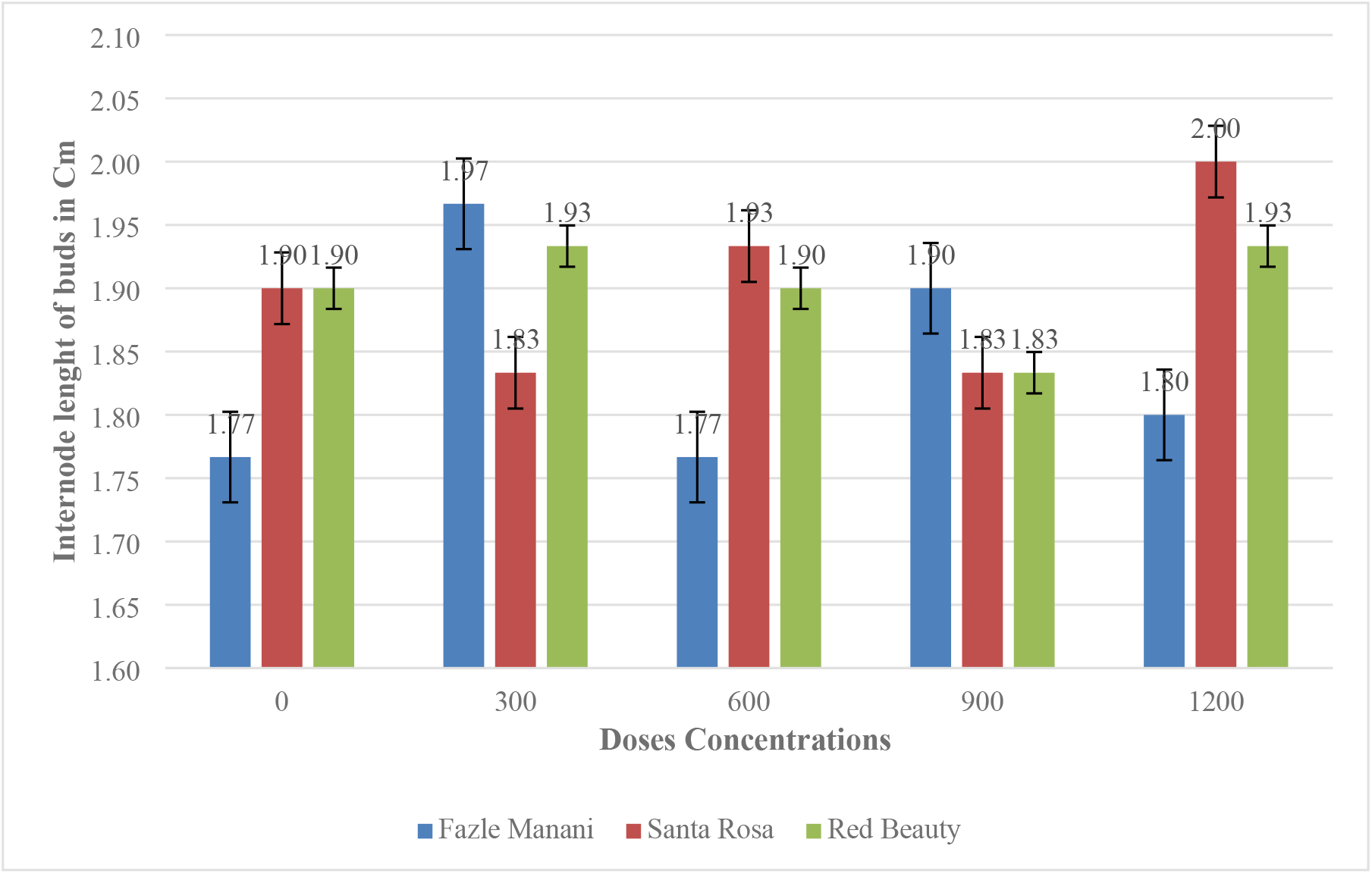
Responses of plum cultivar to various concentrations of IAA in respect of internode bud length

In plum cultivar Santa rosa the highest internode length (2.0 cm) was observed at 1200 concentrations. The lowest inter node length (1.83 cm) was observed at 300 and 900 concentrations. The 600 ppm concentrations also have slightly stimulatory effects on the internode length revealed is (1.93 cm).

The highest internode length (1.93 cm) was recorded in plum cultivar red beauty at 300 and 1200 concentrations. The lowest internode length 1.83 cm was revealed at 900 concentrations. The 600ppmconcentrations also have slightly stimulatory effect on internode bud length revealed is (1.90 cm).

## 4. Discussion

The effect of different concentrations of IAA on days to sprouting,% sprouting, plant height, internode length, No of leaves and number of branches per buddling of three cultivars of plum Fazli manani, Santa rosa and red beauty were carried out. The data revealed that Fazle manani exhibited the shortest days to sprouting, indicating its higher sensitivity to the IAA. Santa Rosa followed a similar trend, although with slightly more number of days to sprouting. Red Beauty, on the other hand, showed the highest days to sprouting, suggesting a lower sensitivity to IAA. It was also observed that higher dose of IAA resulted in lowest days to sprouting except in cultivar red beauty. It demonstrates that an optimal IAA dose can potentially accelerate the sprouting process. According to Fu *et al*. (2015) IAA promotes cell division and elongation in the cells of the bud, leading to the growth and expansion of tissues. It induces the production of proteins and enzymes that are involved in cell division and expansion, thereby facilitating the initiation and development of new shoots. The current study agrees with the findings of Al-Zebari *et al*. (2016) who reported similar results for apricot varieties treated with IAA.

For plant height, the results indicate that IAA has a significant impact on plant height in plum varieties. Fazle manani has the shortest plant height across all the applied concentrations of IAA except 300 ppm concentrations where it slightly increased than control. It showed that higher concentrations of IAA have inhibitory effect on plant height. On the other hand, Santa Rosa and Red Beauty exhibited variable responses to IAA concentrations. Santa rosa showed a relatively higher response particularly to 300 ppm concentrations where plant height significantly declined while increasing the concentrations of IAA, so it showed that higher concentrations have inhibitory effect on plant height, while Red beauty appeared to have uniform plant height against the tested concentrations of IAA. These variations in the response of plum varieties to IAA concentrations can be attributed to genetic and physiological differences among the cultivars. Genetic factors play a crucial role in determining the sensitivity and responsiveness of plants to growth regulators like IAA (Nakamura *et al*., 2003; Majid *et al*., 2019). The findings of this study agree with previous research conducted by Kako *et al*. (2012) on the effects of IAA and other PGR on plant growth of different varieties of plum who reported almost similar results. IAA, as a naturally occurring auxin, is known to promote cell elongation and division, which ultimately contribute to increased plant height.

For the number of branches, the data provided shows that different concentrations of IAA had different effects on the parameter in three cultivars of plum. Looking at the results, it was observed that the plum varieties responded differently to the varying concentrations of IAA. For example, Fazle manani had the highest average number of branches per budding at all IAA concentrations, while Santa Rosa had the highest number of branches at a medium IAA concentration (300 ppm). Red Beauty, on the other hand, had a relatively consistent number of branches across the tested IAA concentrations. These findings suggest that the response of plum varieties to IAA in terms of branch development is specific to each variety. Some varieties showed more responses to higher concentrations of IAA, while others show optimal development at moderate concentrations. These results confirm the findings of Ali *et al*. (2019) who reveled different concentrations of PGR had both stimulatory and inhibitory effect on number of branches and other growth attributes of plum verities.

Results about the effects of different concentrations of IAA on percent sprouting of buddling on plum varieties showed that Red Beauty has the highest average sprouting percentage, indicating that it produced more sprouts in response to the applied concentrations of IAA. Santa Rosa has the lowest average sprouting percentage, suggesting that it is less affected by the IAA concentrations. Fazle manani falls in between the other two cultivars. Comparing the sprouting percentages among the cultivars at each IAA dose, at the highest dose of 1200, Red Beauty has a sprouting percentage of (61.00%), while Fazle manani and Santa Rosa have sprouting percentages of (48.00%) and (39.00%), respectively. This suggests that Red Beauty was positively affected by IAA. On the other hand, Fazle manani produced higher sprouting percentages as compared to Santa Rosa at the different IAA concentrations. The present results agree with Kaur (2017) who reported that application of PGR enhanced sprouting capacity and vegetative growth of peach. Similar results were also obtained by Al-Zebari *et al*. (2016) who studied the effects of different PGR on sprouting success of two cultivars of apricot.

The study on the number of leaves per budling in different plum varieties revealed that each cultivar responded differently to varying concentrations of IAA. In the Fazle Manani cultivar, an IAA concentration of 900 resulted in the highest number of leaves per budling, indicating a positive response to this concentration. For the Santa Rosa cultivar, IAA concentrations of 300 and 600 led to an increase in leaf numbers, while higher concentrations of 900 and 1200 had a negative effect. In contrast, the Red Beauty cultivar exhibited a mixed response, with the highest number of leaves per budling observed at IAA concentrations of 0 and 1200. These findings disagree with the results of El-Hodairi *et al*. (1999) who demonstrated that different IAA concentrations had no effect on number of leaves in date palm. It is possible that different plant species may show different responses to the PGR. On the other hand, results if this study are in close agreement with Koshita *et al*. (1999) who revealed that IAA and other PGR had positive effects on number of leaves and flower bud formation of mandarin trees (*Citrus unshiu*).

The study on budding diameter in different plum cultivars at various IAA concentrations showed varying results. The effect of IAA on the budding diameter of Fazle Manani was non-significant, with the largest diameter (7.87) observed at 0 IAA concentration. For the Santa Rosa cultivar, the effects were significant, as the budding diameter increased with higher concentrations of IAA. In contrast, the Red Beauty cultivar exhibited a mixed response to different IAA concentrations, but these effects were not statistically significant. Al-Zebari *et al*. (2016) and Kako *et al*. (2020) revealed similar results about the effect of IAA and some other PGR on peach and apricot cultivars which strongly agree with the findings of this study. Moreover, the results also partially agree with the study of Firde *et al*. (2020) which revealed both stimulatory and inhibitory effects on plum varieties.

Data for internode length of three plum varieties showed no significant effects of IAA on the internode length. Fazle manani has an internode length mean of (1.84 cm), while Santa Rosa and Red Beauty have means of (1.90 cm). This indicates that, on average, Santa Rosa and Red Beauty tend to have slightly longer internode lengths compared to Fazle manani. However, the differences in internode length among the cultivars are relatively small *Kodad et al*. (2021) reported that internode of almond treated with IAA and PGR significantly increased which disagree with the findings of this study.

## Conclusion

The present study concluded that the most effective dose of PGRs was a concentration of 300 ppm. These findings demonstrate the differential sensitivity of plum cultivars to various concentrations of IAA and highlight its potential for regulating their growth and morphology.

## Availability of data and material

The authors confirm that the data and materials supporting the findings of this study are available within the article.

## REFERENCES

1. Ali, T. J. M., Alwan, M. I., & Obaid, Z. O. (2019). Effect of spraying of gibberellic acid (GA_3_) and foliar fertilizer (Agro leaf) on the seedling growth of plum cultivar ‘Hollywood’. Research on Crops, 20(2), 313–321.

2. Al-zebari, S. M. K., Garou, S. A. M., & Tawfik, S. I (2016). Impact of growth regulators in budding success percentage and growth of two apricots (Prunus armenica l.) Cultivars. In The 2^nd^ Scientific Agricultural Conference (p. 499).

3. El Hodairi, M. H., Amer, A. A., & El Fagih, A. S. (1991). The effects of indole acetic acid (IAA), indole butyric acid (IBA) and naphthalene acetic acid (NAA) on the growth of Taaghiyaat date palm (Phoenix dactylifera L.). Frontier in Tropical Fruit Research 321, 326–333.

4. Firde, K., Seleshi, G., Mossie, T., Setu, H., & Negash, E. (2020). Response of Two Plum Rootstock Varieties to Different Concentrations of Indol-3-Butyric Acid. J Agri Sci Food Res, 11,p284.

5. Fishel, F. M. (2006). Plant growth regulators. EDIS, 2006(6).

6. Flasiński, M., & Hąc-Wydro, K. (2014). Natural vs synthetic auxin: Studies on the interactions between plant hormones and biological membrane lipids. Environmental research, 133, 123–134.

7. Fu, S. F., Wei, J. Y., Chen, H. W., Liu, Y. Y., Lu, H. Y., & Chou, J. Y. (2015). Indole-3-acetic acid: A widespread physiological code in interactions of fungi with other organisms. Plant signaling & behavior, 10(8), e1048052.

8. GoP. 2009. Agricultural Statistics of Pakistan. District wise Area and Production of Plums. Economic Wing, Islamabad (Pakistan). p. 190–191.

9. Kako, S. M. (2020). Effect of plant growth regulators on budding success percentage and growth of sweet cherry transplant (Prunus avium l.). Plant cell biotechnology and molecular biology, 39–46.

10. Kodad, S., Melhaoui, R., Hano, C., Addi, M., Sahib, N., Elamrani, A., … & Mihamou, A. (2021). Effect of culture media and plant growth regulators on shoot proliferation and rooting of internode explants from Moroccan native Almond (Prunus dulcis Mill.) genotypes. International Journal of Agronomy, 2021, 1–10.

11. Koshita, Y., Takahara, T., Ogata, T., & Goto, A. (1999). Involvement of endogenous plant hormones (IAA, ABA, GAs) in leaves and flower bud formation of satsuma mandarin (Citrus unshiu Marc.). Scientia Horticulturae, 79(3-4), 185–194.

12. Majid, S., Hussain, S., Rehman, S.U., Ahmed, M., Ali, W., & Shah, S.H. (2019). Genetic diversity analysis of different maize hybrids for yield and yield components. Journal of Agricultural Science, 11(5), 1–11.

13. Nakamura, A., Nakajima, N., Goda, H., Shimada, Y., Hayashi, K., & Nozaki, H. (2003). Arabidopsis auxin response factors promote expression of the brassinosteroid receptor gene BRI1. Plant Journal, 34(2), 168–175.

14. Neumüller, M. (2011). Fundamental and applied aspects of plum (Prunus domestica) breeding. Fruit, Vegetable and Cereal Science and Biotechnology, 5(1), 139–156.

15. Ohata, K., Togano, Y., Matsumoto, T., Uchida, Y., Kurahashi, T., & Itamura, H. (2017). Selection of prune (Prunus domestica L.) cultivars suitable for the East Asian Temperate Monsoon Climate: Ripening characteristics and fruit qualities of certain prunes in a warm Southwest Region of Japan. The Horticulture Journal, 86(4), 437–446.

16. Okie, W., Hancock, J. (2008). Plums. In: Hancock, J.F. (eds) Temperate Fruit Crop Breeding. Springer,

17. Small, C. C., & Degenhardt, D. (2018). Plant growth regulators for enhancing re-vegetation success in reclamation: A review. Ecological engineering, 118, 43–51.

18. Topp, B. L., Russell, D. M., Neumüller, M., Dalbó, M. A., & Liu, W. (2012). Plum. Fruit breeding, 571-621. Fruit breeding (pp. 571–621). Springer, Boston, MA.

19. Westwood, M.N., (1993). Temperate Zone Pomology. W.H. Freeman and Company, San Francisco, California, USA, p. 223.

20. Wolella, E. K. (2017). Surface sterilization and in vitro propagation of Prunus domestica L. cv. Stanley using axillary buds as explants. Journal of Biotech Research, 8, 18–25.

21. Zhao, Y. (2010). Auxin biosynthesis and its role in plant development. Annual review of plant biology, 61, 49–64.

